# fastBMA: Scalable Network Inference and Transitive Reduction

**DOI:** 10.1101/099036

**Authors:** Ling-Hong Hung, Kaiyuan Shi, Migao Wu, William Chad Young, Adrian E. Raftery, Ka Yee Yeung

**Author notes:** Corresponding author: Ka Yee Yeung.

## Abstract

**BACKGROUND:** Inferring genetic networks from genome-wide expression data is extremely demanding computationally. We have developed fastBMA, a distributed, parallel and scalable implementation of Bayesian model averaging (BMA) for this purpose. fastBMA also includes a novel and computationally efficient method for eliminating redundant indirect edges in the network.

**FINDINGS:** We evaluated the performance of fastBMA on synthetic data and experimental genome-wide yeast and human datasets. When using a single CPU core, fastBMA is up to 100 times faster than the next fastest method, LASSO, with increased accuracy. It is a memory efficient, parallel and distributed application that scales to human genome wide expression data. A 10,000-gene regulation network can be obtained in a matter of hours using a 32-core cloud cluster.

**CONCLUSIONS:** fastBMA is a significant improvement over its predecessor ScanBMA. It is orders of magnitude faster and more accurate than other fast network inference methods such as LASSO. The improved scalability allows it to calculate networks from genome scale data in a reasonable timeframe. The transitive reduction method can improve accuracy in denser networks. fastBMA is available as code (M.I.T. license) from GitHub (https://github.com/lhhunghimself/fastBMA), as part of the updated networkBMA Bioconductor package (https://www.bioconductor.org/packages/release/bioc/html/networkBMA.html) and as ready-to-deploy Docker images (https://hub.docker.com/r/biodepot/fastbma/).

## Findings

### BACKGROUND

Genetic regulatory networks capture the complex relationships between biological entities which help us to identify putative driver and passenger genes in various diseases [1, 2]. Many approaches have been proposed to infer genetic networks using gene expression data, for example, co-expression networks [3], mutual information-based methods [4, 5], Bayesian networks [6-8], ordinary differential equations [9, 10], regression-based methods [11-14] and ensemble methods [15]. In addition, methods have been proposed to infer gene networks using multiple data sources, e.g. [16-19].

#### Our Contributions

We have previously described ScanBMA [14], an implementation of Bayesian model averaging (BMA) [20] for inferring regulatory networks. ScanBMA is available from the “networkBMA” Bioconductor package [21], written in R and C++. It has been shown that ScanBMA generates compact accurate networks that can incorporate prior knowledge.

In this paper, we present fastBMA, which is completely written in C++, and uses more efficient and scalable regression and hashing methods. The algorithmic improvements increase the speed by a factor of 30 on smaller sets, with greater increases observed on larger sets due to improved scalability. fastBMA is parallelized using both OpenMP and MPI allowing for further increases in speed when using multiple cores and processors. Although fastBMA uses the same core methodology as ScanBMA, the increased scalability allows for more thorough sampling of the search space to increase accuracy. The new probabilistic hashing procedure used by fastBMA is faster and utilizes 100,000 times less memory when analyzing large numbers of variables. This allows fastBMA to operate on genome scale datasets without limiting the possible regulators of a given gene to a smaller subset.

A new feature of fastBMA is the implementation of a novel method for eliminating redundant indirect edges in the network. The post-processing method can also be used separately to eliminate redundant edges from networks inferred by other methods. The code is open-source (M.I.T. license). fastBMA is available from GitHub (https://github.com/lhhunghimself/fastBMA), in R as part of the networkBMA package ((https://www.bioconductor.org/packages/release/bioc/html/networkBMA.html)) and as Docker images (https://hub.docker.com/r/biodepot/fastbma/). The Docker containers include all the supporting dependencies necessary for MPI and make it much easier to run fastBMA on a local or cloud cluster.

### RELATED WORK

#### Bayesian model averaging (BMA)

We can formulate gene network inference as a variable selection problem where the dependent variable (target gene expression) is modeled as a function of a set of predictor variables (regulatory gene expression). A regression model can be formed by fitting (1).

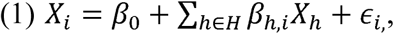

where *X_i_* is the expression of gene i, *H* is the set of regulators for gene i in a candidate model, *β′s* are the regression coefficients, and 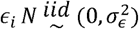 is jhe error term for gene i=1…n.

Time series data can also be modeled by using the expression at the previous time point to predict the next time point.

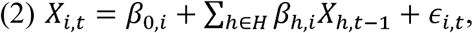

where *X*_i,t_ is the expression of gene i at time t, *H* is the set of regulators for gene i in a candidate model, *β′s* are the regression coefficients, and 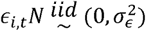 is the error term for gene i=1…n and time t=2,…T.

Different candidate models can be constructed from different sets of regulator genes. Models can be evaluated based upon a measure of their goodness of fit, such as the sum of residuals. However, in genetic analyses, the number of genes often exceeds the number of samples, and many different models can fit the data reasonably well. The core idea behind the BMA methods is to find the posterior probability for each model and make a consensus prediction giving proportionately more weight to the higher confidence models. In terms of gene regulation, the posterior probability that gene *j* is a regulator of gene *i* is the sum of the posterior probabilities of all candidate models that include gene *j* in the set of regulators of *i*. This posterior probability becomes the weight of the edge drawn from gene *j* to gene *i* in the gene network.

#### Estimating model posterior probabilities

Estimation of the posterior probabilities of the models can be accomplished by a variety of methods, some of which are very computationally intensive [12]. The original BMA [20] and iterative BMA (iBMA) methods [22] use the Bayesian Information Criterion (BIC) [23] which is simple to calculate and penalizes larger models which are easier to fit. However, BIC is an asymptotic approximation that is most accurate for large sample sizes. As an alternative, ScanBMA provided the option of using Zellner’s *g* prior [24] to compute the posterior probabilities. The optimal *g* prior parameter is obtained by finding the value that maximizes the total posterior probability of the models. Adjusting the range of possible values for the *g* prior allows us to tune the method for smaller sample sizes and produce better networks. fastBMA exclusively uses the *g* prior to estimate the posterior probabilities and uses faster C++ code for the expectation maximization (EM) optimization of the *g* parameter.

#### Sampling candidate models

The number of possible candidate models grows exponentially with the number of possible regulators, necessitating an efficient methodology to find a subset of reasonable models. In the original implementation of BMA, the leaps and bounds algorithm [25] is used to identify the *n* best models for a given number of variables. Occam’s window [26] is then used to discard models with much lower posterior probabilities than the best model. The leaps and bounds algorithm scales poorly and is limited in practice to fewer than 50 variables. Iterative BMA (iBMA) uses a pre-processing step to rank all variables (genes), iteratively applies the original BMA to the top *w* variables (*w*=30 by default), and discards predictor variables with low posterior inclusion probabilities [13]. In the iterative step, new variables from the ranked list are added to replace the discarded variables. This procedure of repeatedly applying BMA and variable swaps is continued until the *w* top ranked variables have been processed. In contrast to iBMA, ScanBMA removes the restriction of the search space to an initial list of variables [14]. ScanBMA keeps a list of the best current linear regression models found so far and adds or removes a variable from these models to search for better models. The process is repeated until no new models are added or removed from the best set of models. ScanBMA’s greedy approach and the implementation of its core routines in C++ enable it to typically run faster than iBMA. In this paper, we present fastBMA that uses the ScanBMA approach but exploits the fact that new models are based upon existing models. In particular, new models are fitted using the results from the existing models which increases the speed and scalability of the search.

#### Post-processing graphs by transitive reduction

BMA and other methods for reconstructing biological networks can generate edges between genes that are the result of indirect regulation through one or more intermediate genes. While having edges that represent either direct or indirect interactions is a perfectly acceptable graph, biological networks are usually represented by edges that represent direct interactions. Such networks allow for more straightforward identification of potential driver genes. For genetic networks, it is therefore desirable to remove edges between nodes where the regulation is indirect (transitive reduction). This can be done through post-processing of the inferred network. One intuitive approach is based on eliminating direct edges between two nodes when there is a better indirect path [27]. For example, *Bosnacki* recently proposed comparing p-values of the best edge in an indirect path with that of the direct path [28]. fastBMA introduces a similar approach that reduces transitive reduction to a shortest path problem which can be solved more efficiently for the sparse graphs typically found in gene regulatory networks.

Table 1 summarizes the key differences between the different BMA implementation.

**Table 1.**
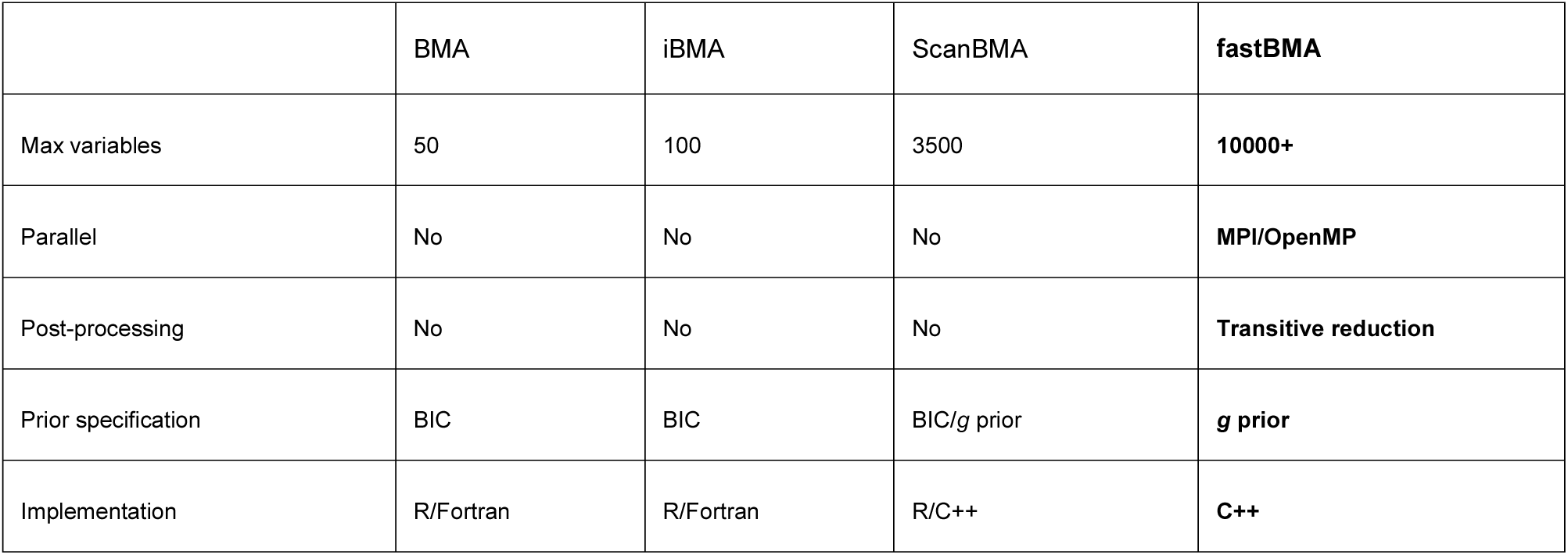
Differences between BMA implementations

|  | BMA | iBMA | ScanBMA | **fastBMA** |
| --- | --- | --- | --- | --- |
| Max variables | 50 | 100 | 3500 | **10000+** |
| Parallel | No | No | No | **MPI/OpenMP** |
| Post-processing | No | No | No | **Transitive reduction** |
| Prior specification | BIC | BIC | BIC/*g* prior | ***g* prior** |
| Implementation | R/Fortran | R/Fortran | R/C++ | **C++** |

### METHODS

Figure 1 shows an outline of fastBMA. In this section, we report our algorithmic and implementation contributions in fastBMA and our evaluation procedure. Pseudocode for the entire implementation is provided in supplemental materials.

**Figure 1.**
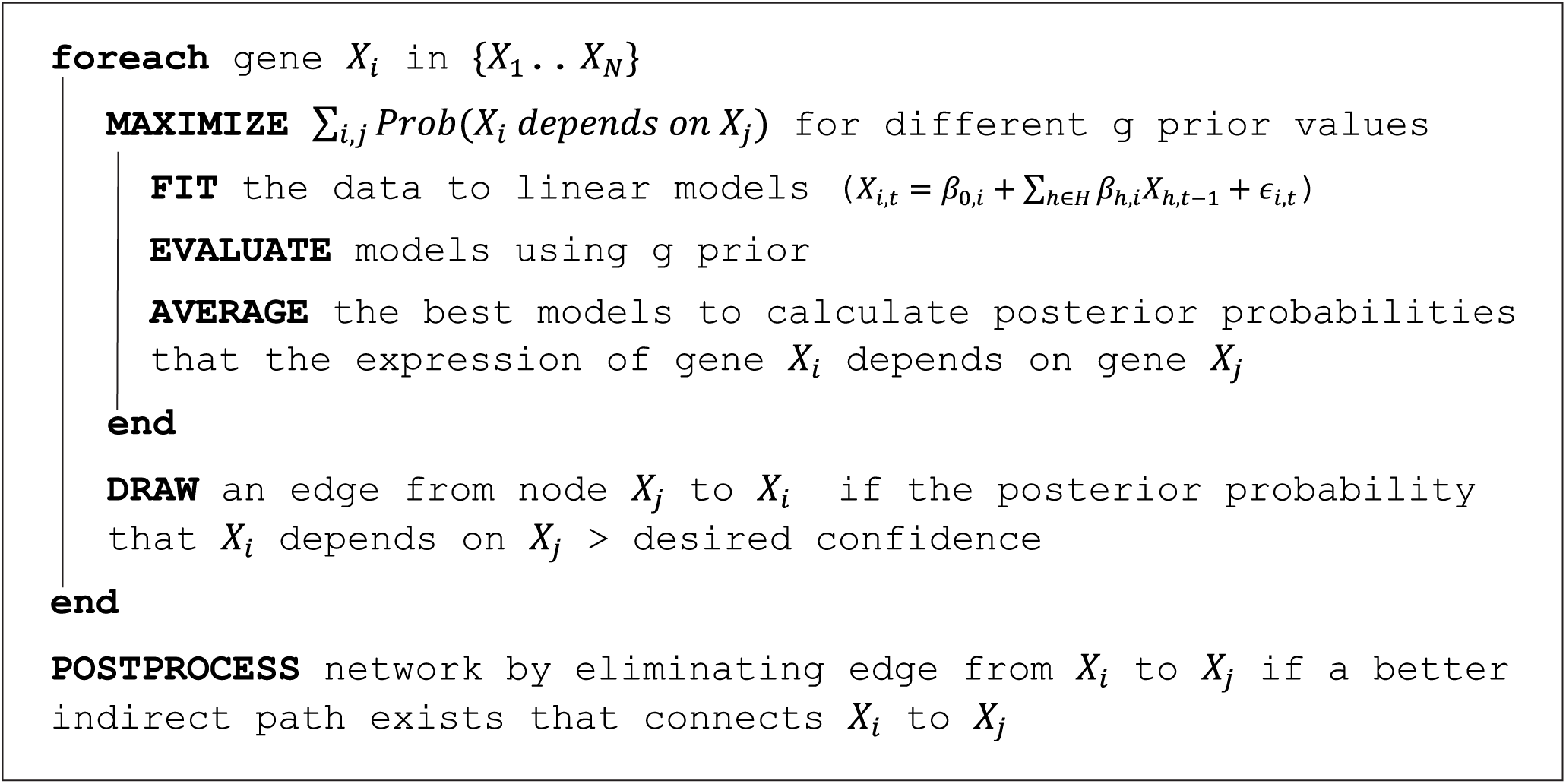
Outline of fastBMA algorithm.

#### Algorithmic outline of fastBMA

The core approach for fastBMA is similar to that used by ScanBMA. The best models are found using ScanBMA’s search strategy with a starting value of *g* in the interval [1… NumberOfSamples]. Brent minimization [29] is then used to find the value *g* in the interval that gives rise to the set of models with the highest marginal probabilities. A graph is constructed by drawing edges between genes with an edge weight equal to the average posterior probability of the regulator over the set of reasonable models. Transitive reduction is applied to this graph to remove edges that can be adequately explained by a better indirect path. A final graph is constructed by retaining edges with weights greater than a given cutoff. There are 4 major algorithmic improvements that increase the speed, scalability and accuracy of fastBMA:

1. Parallel and distributed implementation
2. Faster regression by updating previous solutions
3. Probabilistic hashing
4. Post-processing with transitive reduction

#### Parallel and distributed implementation

Parallelization can be accomplished by using a shared memory system, such as OpenMP (http://openmp.org/wp/), which is designed for assigning work to different threads in a single CPU with multiple cores. In contrast, MPI (Message Passing Interface) (https://www.mpich.org/) launches multiple processes on one or more CPUs and passes messages between processes to coordinate the distribution of work. Both of these approaches have their respective advantages and disadvantages. OpenMP is applicable only to CPU’s on a single machine and is a bit slower for fastBMA. MPI is usable on a single machine or a cluster but requires some work to set up. fastBMA implements both approaches allowing the user to choose the preferred methodology based on their requirements.

Inferring the entire regulatory network involves finding the regulators for every gene in the set. Since each of these determinations is carried out separately, each thread or process can be assigned the task of finding the regulator for a subset of genes in the set. When OpenMP is used, it provides a scheduler that dynamically assigns the regression calculations for a given gene to each thread. Threads work simultaneously on their tasks and receive a new task when they finish the previous task. All threads share access to memory and the same input data for the regression is available to all the threads. The parallel code only extends to the regression loop - the final transitive reduction post-processing and output is done by a single thread.

When MPI is used, we initially split the tasks evenly among the available CPUs. In the case of MPI processes, memory is not shared. Instead the input data is read by a master process and distributed to all the participating processes using MPI’s broadcast command. All processes then work on their tasks simultaneously in parallel and send messages to all the other processes so that all processes know which tasks are being worked upon. The length of time required for each calculation varies considerably and as a result, some processes will finish before others. A process that finishes early then works on tasks initially assigned to other processes that have not yet been started. When all the regulators for all the genes have been found, a master process gathers the predictions, performs transitive reduction post-processing and outputs the final complete network. OpenMP can also be used in conjunction with MPI to further subdivide the tasks among threads available to a CPU.

#### Faster regression by updating previous solutions

Even with the above parallel implementation, each individual calculation of regulators is still accomplished by a single process. If the regression procedure is too slow, this step can be rate-limiting for large numbers of genes regardless of the number of processors available. ScanBMA uses Cholesky decomposition to triangularize the regression matrix and obtain the regression coefficients through back substitution. These calculations have a time complexity of O(*n*^3^) where *n* is the number of variables in the model. However, in the case of fastBMA, new regression models are based upon the previous models and involve the addition or removal of a single variable. It is possible to use the triangular matrix of the previous model to calculate the triangular matrix and regression coefficients for the new model. fastBMA’s new C++ implementation of this update algorithm is based on the Fortran code from the qrupdate library (http://sourceforge.net/projects/qrupdate/).

The Cholesky decomposition becomes O(*n*^2^) when updating the previous solution. Average sampled model sizes for typical applications range between 5-20 and this would be the expected speedup when using a single thread. However, fastBMA further optimizes the implementation by pre-calculating matrix multiplications and using lower level linear algebra routines from OpenBLAS (http://www.openblas.net/) for further speed increases. OpenBLAS is an optimized open source implementation of the BLAS (Basic Linear Algebra Subprograms) routines. Custom wrappers were added to allow the use of the OpenBLAS Fortran libraries. The improvements in the regression procedure account for the majority of the 30-fold increase in speed observed for smaller search spaces on a single thread.

#### Replacing the hash table with a constant time and constant space probabilistic filter

Before evaluating a newly generated model, ScanBMA checks to see if that model has been previously evaluated. This is done by using a hash table to store a string representing the indices of the variables in the model. For smaller sets, the time and space required for this operation are negligible compared to the time and space required to calculate the regression coefficients. However, when the number of variables is in the thousands, this operation becomes the bottleneck. A regular hash table uses a hash function to map the model to a bucket. When the number of models is small relative to the number of buckets (small load factor) it is unlikely that two models will be put in the same bucket and the time taken to look up a model is just the time to map the model to a bucket. For lexicographical strings, the hash function is applied to small substrings and the values are combined. The time required for hashing the whole string is proportional the length of the string. In the case of ScanBMA, the length of the strings formed from the concatenated variable indexes is proportional to the number of variables *n*. Thus for small numbers of models, the time complexity of the lookup operation will also be O(*n*),

However, when the load factor is large, it is likely that multiple models map to the same bucket. The resulting collisions must be resolved by searching through the models in the bucket. For the C++ *unordered set* container used by ScanBMA, this has worse case O(*m*) time complexity where *m* is the number of models giving a total time complexity of O(*nm*) for the lookup procedure when *m* is large. In addition, the memory required to store the hash table will be O(*m*). Unfortunately, when a large number of mostly uninformative variables are coupled with a large Occam’s window, *m* grows very rapidly. In these cases, we observed that the memory and time requirements of the hashing procedure soon become limiting. For example, even though it only runs a single thread, ScanBMA will crash a 56 GB machine when there are large numbers of variables and no informative priors.

It is vital that the ScanBMA algorithm does not sample a model more than once to ensure that the method will converge and terminate. However, the methodology is quite tolerant of falsely excluding models that have not been sampled. ScanBMA only explores a small sample of the possible models – the vast majority of models are normally excluded. Furthermore, in the BMA approach, many models are averaged to obtain the final edges. Variables that are important appear in many models. In the rare case where a good model is falsely excluded, the impact is minimized because the key regulators in the falsely excluded model will be found in other models. When such false negatives are tolerated, an alternative to using a hash table is to ignore the collisions. This saves both time and space by removing the dependence on *m* for both time and space complexity. An example of a noisy or probabilistic hashing approach is the Bloom filter [30], which has been used for bioinformatics applications [31] due to fast computation and low memory requirements.

fastBMA includes an optimized implementation of a probabilistic hash (see Figure 2) that has constant time and constant memory complexity. The dependence of the computation time on *m* is eliminated by ignoring collisions and the dependence on *n* is eliminated by using an updatable hash function (MurmurHash3: https://github.com/aappleby/smhasher) that calculates the hash value of a model based on the hash value of the previous model. fastBMA uses the hash value of the model to map it to a location in a two dimensional bit table. The bit at that location is then set to 1. Any model that hashes to a table location with a set bit will not be processed. The error rate for the filter is initially very low and errors are more likely near the end of the search when more bits in the table have been set. This meshes well with the search process used by fastBMA: errors at the end of the search have even less impact because almost all changes to good models are rejected at that point.

**Figure 2.**
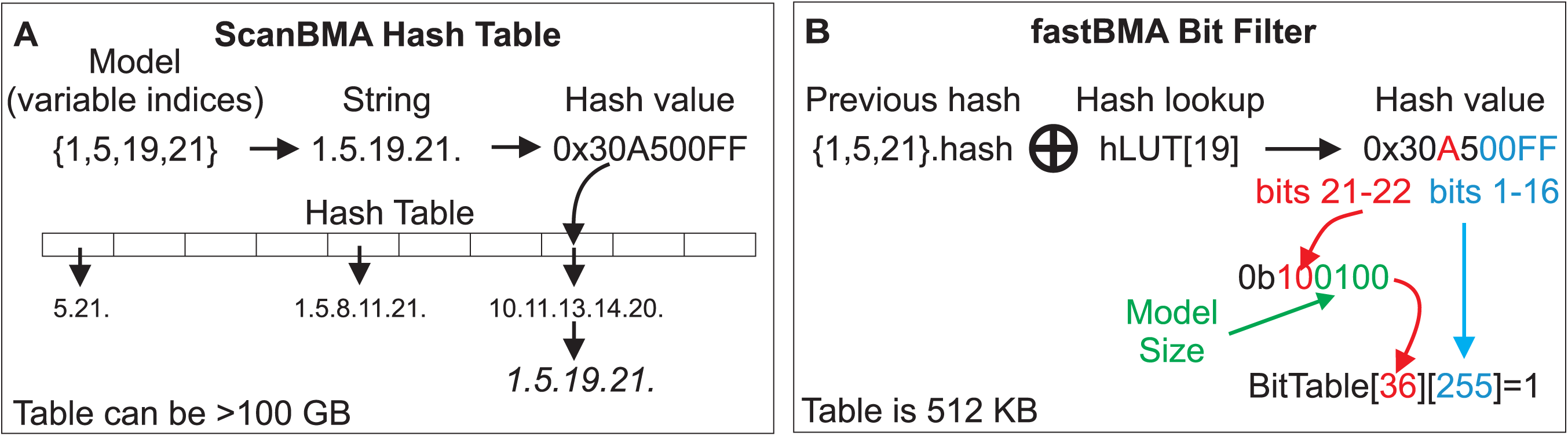
ScanBMA hash table versus fastBMA bit filter. The differences between the hashing methods used by ScanBMA and fastBMA are shown. ScanBMA concatenates the indices of the regulator variables in the model to form a unique string. The string is then mapped to one of a set of buckets. Strings mapping to the same bucket are kept in a second data structure that must be navigated to look up the string. In contrast, fastBMA pre-calculates the hashes for all the possible variables. New regression models are based upon the previous models and involve the addition or removal of a single variable. The hash value for the new model is obtained by XORing the hash value for the variable to be added or deleted with the hash value of the previous model. The hash value is used to map the model to a position in a 512 kB bit table with the row dependent on the number of variables. Mapping different sized models to different rows prevents the large number of collisions that would otherwise arise when using the XOR operator to combine hash values. A bit is set in the bit table to indicate that the model has been observed. Collisions are ignored - it is possible to falsely conclude that a novel model has been evaluated when it has not. As discussed in the Methods section, this type of error is well-tolerated by the fastBMA protocol.

**Figure 3.**
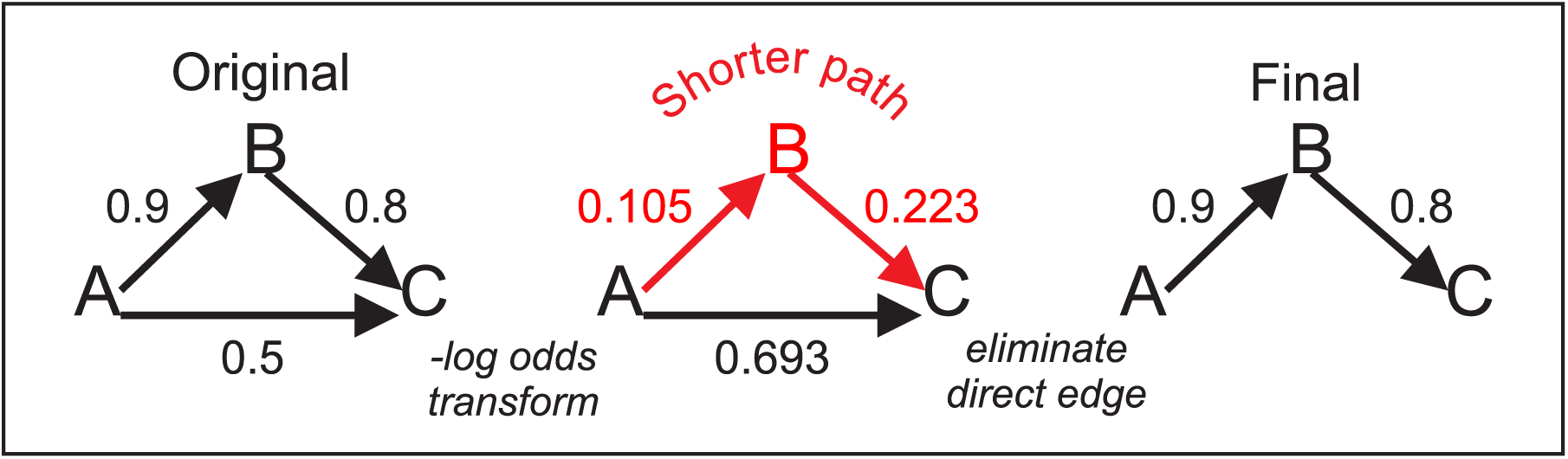
Transitive reduction post-processing. A simple example of the transitive reduction procedure is illustrated. The three edge weights in the mini-graph are the posterior probabilities that A regulates B, B regulates C and A regulates C. The probability of A regulating C through an indirect path through edges A→B→C is the product of the edge weights for A→B and B→C. We take the negative log of the probabilities (middle panel) to transform the multiplication into distances. The indirect path A→B→C is shorter than the direct path A→C which is equivalent to the probability of A regulating C through B being greater than the probability of A directly regulating C. As a result, the edge between A and C is removed.

Our benchmarking confirms that ignoring collisions does not degrade the accuracy of fastBMA. Using a bit table of just 512 **<u>*kilo*</u>**bytes gives identical results for smaller synthetic dataset and almost identical results for the larger genome-wide experimental dataset This is reflected in figure 4 where the accuracy of fastBMA is the essentially the same (actually slightly higher) than ScanBMA when using the same search window. However, ScanBMA can use hundreds of **<u>*giga*</u>**bytes of memory to maintain a string hash table during wide searches over the yeast dataset.

**Figure 4.**
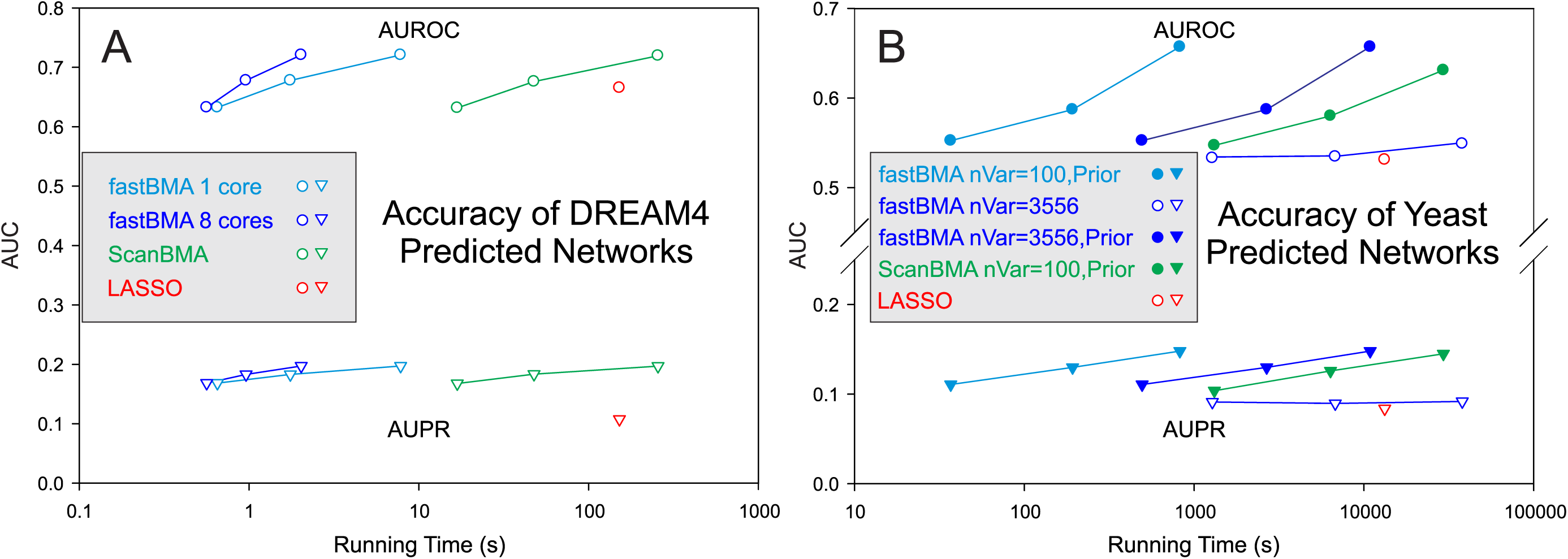
Graphs of the overall accuracy of networks as a function of running time on the DREAM4 simulated and yeast time series data. The area under the receiver operating character curve (AUROC) and area under the precision recall curve (AUPR) of networks inferred from the DREAM4 dataset using fastBMA (no post-processing), ScanBMA and LASSO are plotted against the running times. The different points within a line segment represent fastBMA and ScanBMA with increasingly wider searches as determined by the odds ratio (OR) parameter (OR=100,1000,10000) – the leftmost point representing the smallest OR which is the fastest and least accurate. LASSO does not have an equivalent parameter and was run with the default settings. For the yeast datasets, prior probabilities of regulatory relationships (informative priors) were obtained using external data sources as described in Lo *et al*. For all methods not using informative priors (including LASSO) variables were ordered by their absolute correlation to the response variable. For the ScanBMA on the yeast dataset, the search space was restricted to the 100 variables with the highest prior probabilities. fastBMA was run with a search space of 100 variables using 1 core and all 3556 variables using 8 cores, with and without the Lo et al. prior probabilities. All tests were conducted on an A10 Microsoft Azure cloud instance, which is an Intel Xeon CPU with 8 cores and 56 GB of RAM.

The implementation of the methodology is also further optimized for speed. New hash values are derived from old ones by looking up the value of the pre-calculated hash for the variable to be added or deleted and using XOR to combine it with the previous hash. This procedure is very fast and invertible but normally would cause very severe collision problems with the same hash being associated with different sets of variables. This is solved by mapping hashes from models of different sizes to different rows of the bit table. fastBMA uses a bit table of 64 rows by 65326 columns. fastBMA maps the lower 16 bits of the hash value to obtain the column *c* and uses bits 21 and 22 combined with the last 4 bits of the model size to obtain the row *r* (see Figure 2). The value of the bit table at row *r* and column *c* is set to indicate that the hash value has been seen. Thus the hashing/insert/lookup procedure is constant time, using a very small number of fast bit operations. The tiny size of the bit table (512 kB) also makes the lookup operation very cache friendly. Due to these optimizations, the bit filter used by fastBMA is much faster than using a full hash table even for small datasets where the load factor is small and there are few collisions.

#### Transitive reduction to eliminate redundant edges

fastBMA’s transitive reduction methodology is based on eliminating direct edges between two nodes when there is a better alternative indirect path [27]. *Bosnacki* recently proposed comparing p-values of the best edge in an indirect path with that of the direct path [28]. fastBMA uses the stronger criterion of comparing the overall posterior probability of the entire path. The linear regression model underlying BMA does not distinguish between direct and indirect paths. However, BMA is usually seeded with the prior probabilities of a direct interaction between genes, and the posterior probabilities that constitute the edge weights in a fastBMA network are intended to be estimates of the confidence that there is a direct interaction. The overall probability of any path can be estimated (assuming independence) by multiplying the edge weights together. Equivalently, we can transform the edge weights by taking the negative log and the highest probability path becomes the path with lowest sum of negative log edge weights. The question of whether a better indirect regulatory chain exists is thus mapped to the question of whether a shorter indirect path exists between the two nodes. This is the a shortest path problem that can be solved by Dijkstra’s method with time complexity of O(N E logN + N^2^ logN) where E is the number of edges and N is the number of nodes. By comparison, the GPU methodology of *Bosnacki* is O(N^3^) using a less selective criterion of comparing best edge in the path. The search is also bounded: once a path's distance exceeds the direct distance, there is no need to further explore that path. In addition, fastBMA produces graphs with few high weight edges and in practice, the algorithm is much faster than the worst case as most searches are quickly terminated.

#### Datasets used for testing

We have previously benchmarked ScanBMA[14] against other network inference methods (MRNET [5], CLR[32], ARACNE [4], DBN [8], and LASSO [11, 33]) on smaller test sets. In this study we compare fastBMA only to ScanBMA and LASSO which were the two most accurate methods in these benchmarks and are the only two methods that could infer networks from the larger datasets in a reasonable time.

We used the following 3 datasets for testing.

1. *Simulated* 10-gene and 100-gene time-series data (5 sets of each) and the corresponding reference networks from DREAM4 [34-38]. As these datasets are simulated, the true regulatory relationships are known and are used to evaluate the accuracy of the predicted networks.
2. Yeast time-series expression data (ArrayExpress E-MTAB-412) consisting of 3556 genes over 6 time points [39]. Being actual data, there is no absolute ground truth. Instead, we compared the regulatory predictions with the literature-curated regulatory relationships from the YEASTRACT database [40].
3. Human single-cell time-series RNA-Seq data GSE52529 (9776 genes) from GEO [41]. As no satisfactory gold standard was available, we only used this to demonstrate that fastBMA could scale to noisy human genome-wide expression data.

#### Assessment metrics

We define a true positive (TP) as an edge in the inferred network that is also present in the ground truth or gold standard set. False positives (FP) are edges in the inferred network that are missing in the gold standard. False negatives (FN) are missing edges in the inferred network that are present in the gold standard and true negatives (TN) are missing edges that are also missing in the gold standard. Precision (TP/(TP+FP)) and recall (TP)/(TP+FN) are useful measures of the positive predictive value and sensitivity of the methodology. However precision and recall are dependent on the threshold used for the edge weights. Plots of precision versus recall over different values for the threshold give a more complete picture of the accuracy of the network inference. Similarly, receiver operating characteristic plots of TP/(TP+FN) versus FP/(FP+TN) for different thresholds are also useful, though less so than precision-recall plots because we are more interested in TP in sparse biological networks. We distill the overall information of these plots into a single number by estimating the area under the curve (AUC) i.e. area under precision recall curve (AUPRC) and area under receiver operating curve and (AUROC) for all possible threshold values. Due to the size of the larger yeast networks, all AUC calculations were done using custom software fastROCPRC (https://github.com/lhhunghimself/fastROCPRC) written in C++. We primarily use AUPR and AUROC for the assessment as these metrics measure the overall performance of the methods. In practice, however, predicting *some* edges accurately, even if only for the most confident predictions is still valuable for narrowing down a set of potential interactions to be further explored. Hence, we also plot the precision-recall graph to assess where the differences in accuracy are occurring.

### RESULTS

We applied our fastBMA algorithm to both simulated and real time series gene expression data. We had previously tested several methods on these datasets [14] and found that ScanBMA and LASSO were the fastest and most accurate methods. Therefore we compared the fastBMA results to ScanBMA and LASSO [33, 42]. LASSO is a non-Bayesian linear regression method that uses a penalty term to prevent overfitting to models with many variables. It is written in Fortran and is one of the fastest network inference methods available. Both fastBMA and ScanBMA control the breadth of the search by varying the odds ratio threshold that defines the size of Occam’s window. The odds ratio is the confidence in the query model relative to the best model. Models outside of this window are discarded. Hence, a larger odds ratio threshold drives a wider search which naturally takes longer to complete.

We ran both ScanBMA and fastBMA with increasing larger windows and the time and accuracy as measured by AUROC and AUPR plotted as line segments in figure 4. All the line segments have a positive slope, indicating that larger windows do increase the accuracy, at the expense of using more computation time. For both the synthetic DREAM4 and experimental yeast datasets, with or without prior information, the line segments for fastBMA are well to the left of the corresponding line segments for ScanBMA. The x-axis is logarithmic, indicating fastBMA is orders of magnitude faster than ScanBMA when using the same parameters. Alternatively, one can use a larger odds ratio with fastBMA and obtain a more accurate result in same time it would take to run ScanBMA with a smaller odds ratio. On the same datasets, fastBMA is also more accurate and faster than LASSO, the degree and nature of improvement dependent on whether the user chooses to emphasize speed or accuracy through the choice of the odds ratio parameter.

One of the main advantages of the BMA methods is that they are able to incorporate prior information to improve inference. This was not possible for the DREAM4 dataset as it is a synthetic dataset. In this case, an uninformative uniform prior probability is used. However, for the yeast dataset we had access to priors from external data sources [12]. This also allowed us to triage the variables to be explored to the 100 variables with the highest prior probabilities, saving considerable computational resources. As expected, using informative priors increases the accuracy and decreases the running time relative to LASSO. In addition, we ran fastBMA, without informative priors and without restricting the number of variables (i.e. using all 3556). This is beyond the capabilities of ScanBMA when using wider search windows. Even on this computationally demanding task, inferring the yeast network without informative priors, fastBMA is faster than LASSO with increased accuracy as assessed by AUROC and AUPR.

A common use for computational network inference is to identify a small set of potential regulators that could be verified with further experiments. For this use case, an improvement in the precision of the most confident predictions is more important than a small improvement in the overall performance of the method. As some of the differences in AUC for the yeast dataset are relatively small, we plotted the precision recall curves in figure 5. We see that the precision of the most confident predictions (i.e. lowest recall) is increased. The advantage of using informative priors when available is very clear. However, even when prior knowledge is not available, the fastBMA algorithm is superior which is especially evident in the case of the Dream4 dataset.

**Figure 5.**
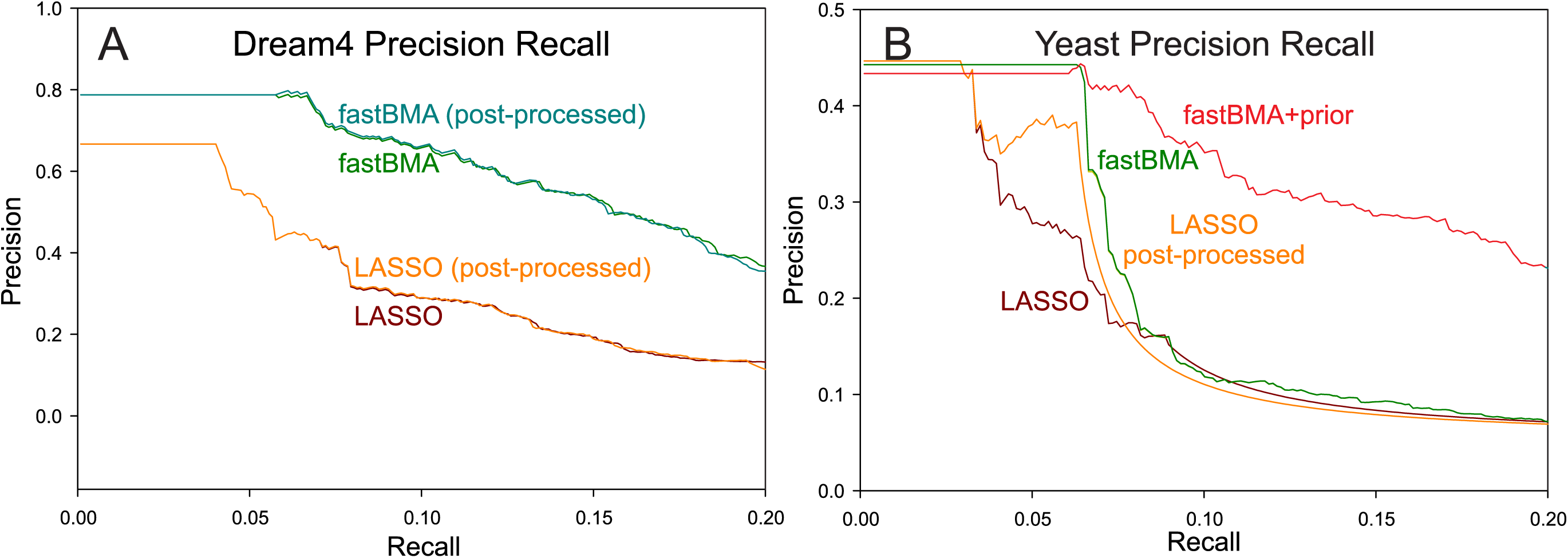
Precision-recall curves. The precision-recall curves were plotted for the networks inferred from the Dream4 data using LASSO, LASSO+post-processing, fastBMA+post-processing. No informative priors were available for this synthetic dataset. Curves were plotted for the networks inferred from the yeast time series data using LASSO, LASSO+postprocessing, fastBMA and fastBMA with informative prior. For the yeast dataset, curves for post-processed networks for fastBMA are not shown as they are essentially identical to the curves for networks inferred without post-processing.

The effect of post-processing is more limited. In figure 5, the precision-recall curves for the Dream4 dataset are almost identical for fastBMA and LASSO with and without post-processing. The same result was observed for fastBMA on the yeast dataset and for clarity, we did not plot the overlapping precision-recall curves for the post-processed networks for fastBMA. However, we do see that postprocessing has an effect on LASSO for the yeast dataset.

We also tested fastBMA on a human single cell RNA-Seq dataset with 9776 variables. Using a 32 core cluster on Microsoft Azure, fastBMA was able to obtain a network in 13 hours without using informative priors. Neither ScanBMA, nor LASSO is able to return results for this dataset. We do not have a gold standard for this test – the purpose was to demonstrate that fastBMA could handle a very large and noisy genomic sized dataset and return a network within a reasonable time even in the worst case scenario where the data is noisy and there is no prior information.

### DISCUSSION AND CONCLUSIONS

We have described fastBMA, a parallel, scalable and accurate method for inferring networks from genome wide data. We have shown that fastBMA can produce networks of increased accuracy orders of magnitude faster than other fast methods even when using a single thread. Further speed increases are possible by using more threads or processes. fastBMA is scalable and we have shown that it can be used analyze human genomic expression data even in the most computationally demanding situation of noisy data, no informative priors and considering all genes a possible regulators.

fastBMA includes a novel transitive reduction post-processing methodology for removing redundant edges where the predicted regulatory edge can be better explained by indirect paths. Both fastBMA and LASSO already penalize large models and favor the exclusion of redundant variables. This explains why post-processing has minimal impact on the sparse networks predicted by fastBMA and LASSO. In particular, fastBMA produces very sparse networks which are not improved by further processing on any of the datasets tested. LASSO’s networks are denser. For the small synthetic DREAM4 set, the post-processing still does not improve the network. However, on the larger experimentally derived yeast dataset, spurious edges do appear in the LASSO networks despite the regularization penalty that discourages larger models. Some of these redundant edges are successfully removed by the transitive reduction post-processing, improving the overall accuracy of the network. The transitive reduction module of fastBMA can be applied to any set of edges, and not just those generated by fastBMA or LASSO. It may prove useful as an adjunct to methods and datasets that give rise to denser networks and are more prone to over-predicting edges.

Although we have focused on biological time series data, fastBMA can be applied to rapidly infer relationships from other high dimensional analytics data. Also the fastBMA methodology can be extended for even more demanding applications. For example, multiple bit filters could be used to (i.e. a Bloom filter) to hash larger search spaces. We anticipate that the efficiency of fastBMA will be especially useful for very large datasets on the cloud where usage is metered. For this purpose, we have provided Docker images (https://hub.docker.com/r/biodepot/fastbma/) with to facilitate deployment on local or cloud clusters.

## Availability and requirements

**Project name:** fastBMA

**Project home page:** e.g. https://github.com/lhhunghimself/fastBMA),

**Operating system(s):** Linux (MacOS and Windows support provided through the Docker container ((https://hub.docker.com/r/biodepot/fastbma/) and Bioconductor package (https://www.bioconductor.org/packages/release/bioc/html/networkBMA.html)

**Programming language:** C++

**Other requirements:** gcc version > 4.8, OpenBLAS, mpich2 (if MPI desired) to compile code.

**License:** M.I.T.

**Any restrictions to use by non-academics:** None other than those required by the license

## Availability of supporting data

*Simulated* 10-gene and 100-gene time-series data (5 sets of each) and the corresponding reference networks from DREAM4 was obtained from https://www.synapse.org/#!Synapse:syn2825304/wiki/71131 Yeast time-series expression data (ArrayExpress E-MTAB-412) consisting of 3556 genes over 6 time points [39] and literature-curated regulatory relationships from the YEASTRACT database [40]. Human time-series RNA-Seq data GSE52529 (9776 genes) were obtained from GEO [41].

## LIST OF ABBREVIATIONS

AUC: area under the curve
AUROC: area under receiver operator curve
AUPR: area under precision recall
BMA: Bayesian model averaging
iBMA: iterative Bayesian model averaging
BIC: Bayesian information criterion
EM: Estimation Maximization

## Declarations

### ETHICS APPROVAL AND CONSENT TO PARTICIPATE

Not applicable.

### CONSENT FOR PUBLICATION

Not applicable.

### AUTHORS CONTRIBUTIONS

LHH conceived and implemented fastBMA. LHH, KS, MW and WCY tested and benchmarked the software. KS and LHH added fastBMA to the networkBMA R package. LHH generated the Docker packages. All authors read and approved the manuscript.

## ACKNOWLEDGEMENTS

We would like to thank Dr. Roger Bumgarner for helpful discussions.

